# The Almond Expression Atlas: A comprehensive multi-tissue transcriptomic resource for almond gene expression

**DOI:** 10.64898/2026.10.30.763019

**Authors:** Eva M. López-Fernández, MA Zwieniecki, A. Tixier, Juan A. Botía, Pedro J. Martínez-García

## Abstract

**Background:** Improving crop adaptation and enhancing agronomic and quality traits requires a deeper understanding of gene expression regulation, for which expression atlases are valuable tools. Almond (*Prunus dulcis* (Mill.) D.A. Webb) is one of the most economically important tree nut crops worldwide. However, an integrated transcriptomic resource for this species is still lacking. Available data remain fragmented across independent studies, limiting their comparability and reuse.

**Results:** Here, we present the first almond gene expression atlas: a comprehensive multi-tissue transcriptomic resource integrating publicly available RNA sequencing data. A total of 142 high-quality untreated and control samples from 17 independent projects were curated through a metadata standardization process and analyzed using a unified bioinformatics pipeline. Standardized expression profiles were generated for 25,225 annotated genes in the *Prunus dulcis* ‘Texas’ v2.0 reference genome. The atlas integrates two complementary expression approaches: a median TPM approach capturing quantitative expression levels and a rank-based strategy implemented using the Bgee expression calling method, designed to improve comparability across heterogeneous datasets. This design enables the identification of tissue-specific candidate genes and the exploration of expression patterns across tissues and developmental stages. Its applicability is demonstrated through two use cases, highlighting its ability to identify biologically meaningful gene sets associated with bud dormancy transitions and to prioritize candidate reference genes for RT-qPCR normalization in almond.

**Conclusions:** The Almond Expression Atlas v1.0 is publicly available through FruitDBase (fruitdbase.csic.es), providing an interactive platform for gene expression exploration and visualization. This resource supports functional genomics studies and facilitates candidate gene discovery in almond, while ensuring data traceability and promoting data reusability.

## Background

Almond (*Prunus dulcis* (Mill.) D.A. Webb) is a major tree nut crop worldwide, with global production exceeding 1.47 million tons annually [1]. Taxonomically, almonds belong to the genus *Prunus*, together with species such as peach (*Prunus persica*), apricot (*Prunus armeniaca*), and plum (*Prunus domestica*), within the family Rosaceae. Almond kernels have diverse industrial applications, and each application demands specific kernel characteristics in terms of size, flavor, appearance, or other organoleptic qualities [2]. In addition, the increasing frequency of extreme climate events, such as heatwaves and extreme droughts associated with climate change, affects the phenological stages of almond trees and, consequently, their productivity [3,4]. This highlights the need to better understand the biological mechanisms and molecular processes underlying yield, responses to adverse conditions, development, and quality traits. This knowledge is essential for developing new breeding strategies that address current needs [5].

Over the last decades, the rapid development of high-throughput technologies, such as next-generation sequencing, has enabled the generation of massive amounts of data, driven in part by a marked reduction in the cost per sample [6,7]. This has improved our understanding of the molecular processes involved in agronomic traits in almond [8]. At the same time, the growing volume of available biological data must be accompanied by appropriate infrastructure for data storage, management, and accessibility. This requirement has motivated the establishment of public repositories such as the European Nucleotide Archive (ENA) [9], Ensembl [10], and the Genome Database for Rosaceae (GDR) [11], which are essential for implementing FAIR data principles [12] through data storage, curation, and accessibility. By centralizing biological datasets, these repositories improve data traceability, supporting reproducibility across studies and enhancing reuse in new research contexts. These advances have also expanded our knowledge of the almond genome. In recent years, several reference genomes have been published, including the initial assembly of the ‘Lauranne’ cultivar [8], followed by improved versions with enhanced annotation and assembly quality for the ‘Texas’ and ‘Nonpareil’ cultivars [13–15], and the forthcoming release of the ‘Carmel’ genome assembly (M. Davis, C. Lee, W. Hazzard, P. Martínez-García, J.G. Monroe, P. Brown, personal communication). These genomic resources have enabled the identification of candidate genes associated with agronomic traits and facilitated marker-assisted selection. While reference genomes provide the sequence and structure of genes, transcriptomic resources are essential to characterize gene activity across tissues, developmental stages and environmental conditions, thereby facilitating the functional interpretation of genomic information.

High-throughput sequencing technologies have also led to an ever-growing number of transcriptomic and other omics datasets. This wealth of data has promoted the creation of integrated resources capable of bringing together datasets generated across independent experiments using harmonized analytical workflows [16]. Among these integrative resources, expression atlases have emerged as valuable platforms for exploring gene expression patterns and have become increasingly widespread in plant biology, as reflected by the availability of openly accessible resources such as the Olive, Cannabis, and Soybean Expression Atlases [17–19]. Despite the increasing number of transcriptomic studies available for almond, expression data remain fragmented across independent projects, limiting their accessibility, comparability and reuse. Consequently, a comprehensive and standardized transcriptomic resource for almond is still lacking.

To address this gap, we developed the Almond Expression Atlas v1.0, the first comprehensive transcriptomic atlas for this species, integrating two complementary approaches to explore the available gene expression data. It is hosted in FruitDBase (fruitdbase.csic.es), a new web repository designed to store transcriptomic and genomic data. By providing a unified and reusable resource, the atlas is expected to accelerate candidate gene discovery and facilitate future transcriptomic research in *Prunus*.

## Materials and Methods

All bioinformatics analyses were performed on the CSIC high-performance computing infrastructure using standard compute nodes equipped with two Intel Xeon Gold 6248R processors (48 cores in total, 3.0 GHz) and 192 GB of RAM. The analyses were implemented using Shell and R (v4.3.1) scripts. R was chosen for its broad use in biostatistics and transcriptomic analyses. Finally, the pipeline, summarized in Figure 1, was implemented in Snakemake (v7.32.3) [20], running under Python (v3.12.3) to ensure reproducibility and portability to other species. The complete Snakemake pipeline, along with the Shell and R scripts, is available at [21].

**Figure 1.**
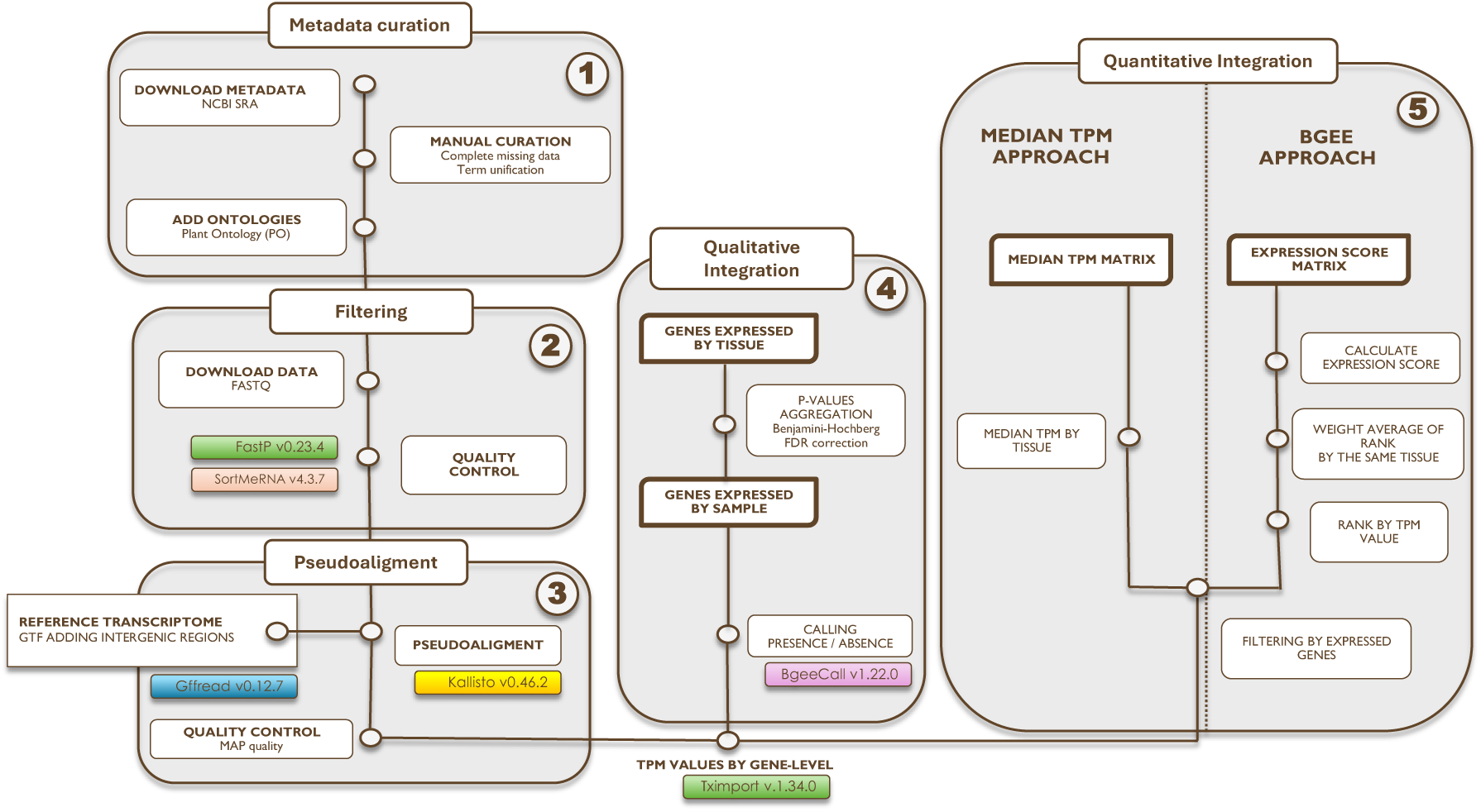
Overview of the RNA-Seq processing and analysis pipeline used for the construction of the Almond Expression Atlas.

### Data collection

To construct a comprehensive gene expression atlas for *Prunus dulcis*, all publicly available RNA sequencing (RNA-Seq) samples for this species deposited in the Sequence Read Archive (SRA) [22] were collected. Only paired-end libraries were selected to standardize processing and minimize technical heterogeneity across samples. SRA is part of the International Nucleotide Sequence Database Collaboration (INSDC) consortium, along with the European Nucleotide Archive (ENA) and the DNA Data Bank of Japan (DDBJ); this repository facilitates programmatic access to samples and their automated processing. Metadata were obtained with the create_sample_info() function of the R package *‘bears’* [17], using the query "*Prunus dulcis*[ORGN] AND rna-seq[STRA]". Following this, a manual metadata curation process was performed to ensure data consistency and to filter out samples with incomplete metadata (see Supplementary Material S1 for the criteria applied). After manual curation was completed, ontology terms were manually assigned to the tissue and developmental stage fields to facilitate data integration and interpretation, ensuring interoperability with other databases and analysis tools and aligning with FAIR principles [12]. Terms were preferentially sourced from the Plant Ontology (PO) [23], which collects anatomical, morphological, and developmental stage terms for plants, and supplemented with the Phenotype And Trait Ontology (PATO) [24] when appropriate terms were not available in PO.

### Filtering and processing data

After metadata curation, raw FASTQ files were downloaded and processed with a custom pipeline adapted from the ‘bears’ package [17]. Using FastP (v0.23.4) [25], reads were filtered for low-quality bases (Phred scores below 20), and adapters were automatically trimmed, generating HTML and JSON reports for each sample. Afterwards, samples with fewer than 5 million reads were discarded, as this is a commonly recommended minimum for gene expression analysis [26]. Additionally, samples with an insert size peak below 100 base pairs (bp) were excluded, as determined by inspection of the insert size distribution across samples (Supplementary Figure S1). Insert sizes below the read length indicate poor library quality, consistent with excessive adapter contamination or over-fragmentation [27].

Next, SortMeRNA (v4.3.7) [28] was used to remove rRNA reads with the pre-built SILVA [29] eukaryotic 18S and 28S rRNA reference databases. The reference transcriptome was generated prior to alignment using gffread (v0.12.7) [30] from a custom GTF file combining exonic regions of the 25,225 protein-coding genes annotated in the *Prunus dulcis* ‘Texas’ v2.0 reference genome [13], restricted to the eight assembled chromosomes (Pd01-Pd08), and intergenic regions obtained following the criteria described in the Bgee expression atlas methodology [31]. This methodology was subsequently used as one of the approaches for assessing gene expression. Pseudoalignment of reads to the reference transcriptome was then performed with Kallisto (v0.46.2) [32] using default parameters to obtain expression estimates.

A summary of sequencing quality and pseudoalignment statistics for all retained samples is provided in Supplementary Table S1.

### Downstream analysis

Expression atlases that combine samples from multiple projects commonly rely on median or mean TPM values without cross-study normalization and use dimensionality reduction techniques to facilitate interpretation [17,19]. Here, two complementary approaches were applied to construct the expression matrices. The first approach relied on median TPM values. The second approach was based on the Bgee expression calling methodology for RNA-Seq data, which combines a ranking-based method with presence/absence calling, in which genes are ranked within each sample and ranks are then integrated by condition to allow comparability across studies [31]. Together, these approaches provide complementary representations of expression: median TPM values offer broader coverage while preserving quantitative expression differences, whereas Bgee provides a more conservative representation based on relative expression.

### Presence/Absence calling

To generate presence/absence gene expression calls, we used the BgeeCall package (v1.22.0). Instead of applying arbitrary abundance thresholds, expression calls were inferred through a statistical comparison against a customized set of intergenic reference sequences, provided through the *custom_intergenic_path* slot of the *UserMetadata* object. Transcript abundance estimates were generated independently using Kallisto and summarized at the gene level using the tximport (v.1.34.0) R package [32] with the ‘lengthScaledTPM’ option to account for transcript length differences. To unify technical replicates, the runs of each sample were merged at the FASTQ level. The resulting gene-level abundance estimates were subsequently formatted to be compatible with the *KallistoMetadata* class. We then executed the *generate_presence_absence* function, which performs the final step of the Bgee workflow and computes a p-value for each gene based on the distribution of intergenic reference abundances. Genes with a p-value < 0.05 were considered confidently expressed. To determine expression at the tissue level, p-values were adjusted across biological samples within each tissue using the Benjamini-Hochberg FDR correction. A gene was considered expressed in a tissue if the FDR value was ≤ 0.05 in at least one sample of that tissue, following the Bgee expression call framework [31].

### Generation of the dimensionality reduction plot

The median TPM across tissues was calculated from the values obtained as described in the previous section, after filtering for genes classified as expressed using presence/absence calls. Since most projects cover only a single tissue type, the median TPM provides an estimate of the expression tendency at the tissue level, while the resulting values also implicitly capture study variability. To explore sample expression patterns, a dimensionality reduction method was applied. The ‘scran’ package [33] was used because its variance-modeling option allows the decomposition of variance into biological and technical components. Raw counts were normalized by library size and log-transformed using the ‘scater’ package [34]. The top 5,000 genes with the highest biological variability were selected using the *getTopHVGs()* function. Subsequently, t-distributed stochastic neighbor embedding (t-SNE) was applied to these highly variable genes to visualize the data distribution in a low-dimensional space. To optimize the t-SNE visualization, three perplexity values were evaluated (10, 15, and 30). A perplexity of 30 was selected because it provided the most interpretable separation of samples in the low-dimensional t-SNE representation [17].

### Generation of the expression score matrix

The expression score matrix was then generated based on the gene-level rank computation for RNA-Seq implemented by Bgee [31]. First, fractional rankings were calculated from TPM values for each sample. Then, for each gene and tissue, the weighted mean of its fractional rankings was computed, with the weights determined by the number of unique ranking positions without ties (Eq. 1, described in [31]). Finally, the expression score of a gene in a tissue was calculated from its global weighted rank and the maximum rank observed across genes and tissues. The rank was first inverted so that lower ranks resulted in higher scores. The resulting value was then normalized to the maximum rank observed across all genes and tissues and multiplied by 100 (Eq. 2, described in [31]). The final scores range from 0 to 100.

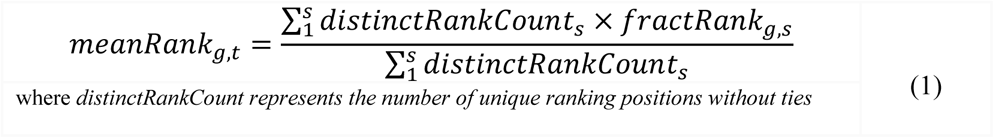

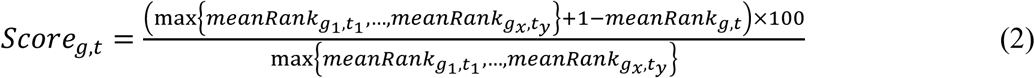

### Identification of expressed and tissue-specific genes

The *Prunus dulcis* ‘Texas’ v2.0 genome assembly was used as the reference [13]. Only genes annotated in Pd01-Pd08 were retained for downstream analyses, excluding unanchored scaffolds and resulting in a final set of 25,225 annotated genes from the 27,969 protein-coding genes reported in the original annotation. To determine whether a gene had detectable expression in a tissue, we applied the same criterion in both approaches, based on presence/absence calling instead of an arbitrary TPM threshold, as described in the previous section and following the Bgee methodology. Tissue-specific genes were defined as those with a Tau index equal to or greater than 0.80 [35]. This index ranges from 0 to 1, where 0 indicates low specificity and 1 indicates high specificity. Tau was calculated separately for each approach: from median TPM values in the median TPM approach and from expression scores in the Bgee approach. For the comparison of tissue-specific gene results between approaches, we calculated the similarity between sets using the Szymkiewicz-Simpson coefficient [36,37], which quantifies shared elements relative to the smaller set.

Presence/absence calls, expression scores, median TPM and tissue specificity were computed in the same way at the developmental-stage level, using tissue-by-developmental-stage combinations instead of tissues as conditions.

### Functional analysis

The functional analysis for tissue-specific genes was performed with the clusterProfiler package (v.4.14.6) [38]. Almond genes were mapped to their *Arabidopsis thaliana* orthologs (TAIR10 protein set). *Arabidopsis thaliana* was selected because, as a model organism, it has an extensive and well-characterized functional annotation. Using DIAMOND blastp (v2.0.13) [39], the best *Arabidopsis* ortholog for each almond gene was selected based on the highest bitscore, with an e-value threshold of 1.5e-5. Finally, the terms with the lowest adjusted p-values were visualized. A GO semantic similarity network was computed using the GOSemSim package (v2.32.0) [40] with the Wang method. To manage redundancy, terms were clustered using the Unweighted Pair Group Method with Arithmetic Mean (UPGMA) with a height cutoff of 0.70. For visualization purposes, each cluster was annotated with the GO term with the lowest adjusted p-value and visualized using a force-directed layout implemented in the igraph (v.2.1.4) R package [41] to facilitate tissue-specific interpretation.

Enrichment analysis was also performed directly on *Prunus dulcis* annotations using STRING (v12.0) [42]. A custom proteome was generated by uploading the *Prunus dulcis* protein sequences (obtained from GDR [11]) to STRING. The resulting organism identifier (STRG0A95YGD) was then used to query the STRING API and perform functional enrichment on each set of tissue-specific genes, retaining terms with FDR < 0.05. Concordance between both analyses was assessed by computing the semantic similarity between their enriched GO term sets again using GOSemSim (v2.32.0) with the Wang method.

### Use cases

For the first use case, genes specific to flower buds at stage T2 were retrieved from FruitDBase, and their GO enrichment was analyzed using STRING through the external link available in the Expression Explorer Bgee-approach view. The EcoGS (Eco Cluster Genes Candidates) proposed by Prudencio et al. [43] were mapped to the *P. dulcis* ‘Texas’ v2.0 annotation by BLAST. A candidate was considered a match when it was classified as specific to flower bud T2 in the atlas.

In the second use case, the median TPM expression matrix by tissue was downloaded from the repository. Traditional and novel reference genes validated by RT-qPCR in *Arabidopsis* by Czechowski et al. [44] were selected (Supplementary Table S7), excluding three hypothetical genes for which no amplification was obtained in the original study. For genes measured with non-gene-specific ATH1 probe sets (EF-1α, 247644_s_at; UBC9, 253826_s_at), all *Arabidopsis* genes detected by the probe set were considered. Almond putative orthologs were retrieved from the DIAMOND best-hit mapping against the TAIR10 protein set described above. When several almond genes shared the same *Arabidopsis* best hit, each of them was analysed separately. For each gene, the coefficient of variation (CV) of the median TPM across the 12 tissues was calculated. Expression stability was expressed as the percentile of each gene’s CV relative to all genes with a mean expression of at least 5 TPM (14,793 genes). Lower percentiles indicate more stable expression. An *Arabidopsis* gene was considered supported as a candidate reference gene for almond when at least one of its putative orthologs ranked within the 25% most stable genes of the atlas.

### Web application development

The web platform was implemented using a client-server architecture. The backend was developed in PHP for request handling and database interaction, while a relational MySQL database stores metadata, derived metadata, transcriptomic data and gene annotation data. The frontend was developed using Tailwind CSS and JavaScript, incorporating DataTables.js for interactive tables and D3.js/Chart.js for data visualization. The application is hosted on CSIC institutional servers.

## Results

### Landscape of publicly available transcriptomic data for *Prunus dulcis*

The initial data retrieval performed in September 2026 identified 305 samples from 27 projects, including 269 publicly available samples from 26 projects and 36 samples from one unpublished project provided by UC Davis (Supplementary Table S3). At the time of the query, the retrieved datasets were predominantly from Spain (39.7% of samples), China (37.4%), and the United States (17.7%) (Fig. 2A-B). This distribution broadly reflects the pattern of global almond production (FAOSTAT; Supplementary Table S2), where the USA is the largest producer, followed by Australia and Spain, with Iran ranking sixth and China thirteenth. In terms of tissue distribution, leaf samples were the most abundant (21.0%), followed by fruit (16.4%), pedicel (13.8%), and flower buds (13.4%) (Fig. 2C). In contrast, specialized tissues, such as phloem, seed coat, and root tissues, were underrepresented or absent, highlighting gaps in the current transcriptomic coverage. Regarding cultivar composition, ‘Zhi pi’ accounted for nearly 20% of all samples, followed by ‘Wan Feng’ (13.8%) and ‘Lauranne’ (12.5%) (Fig. 2D).

**Figure 2.**
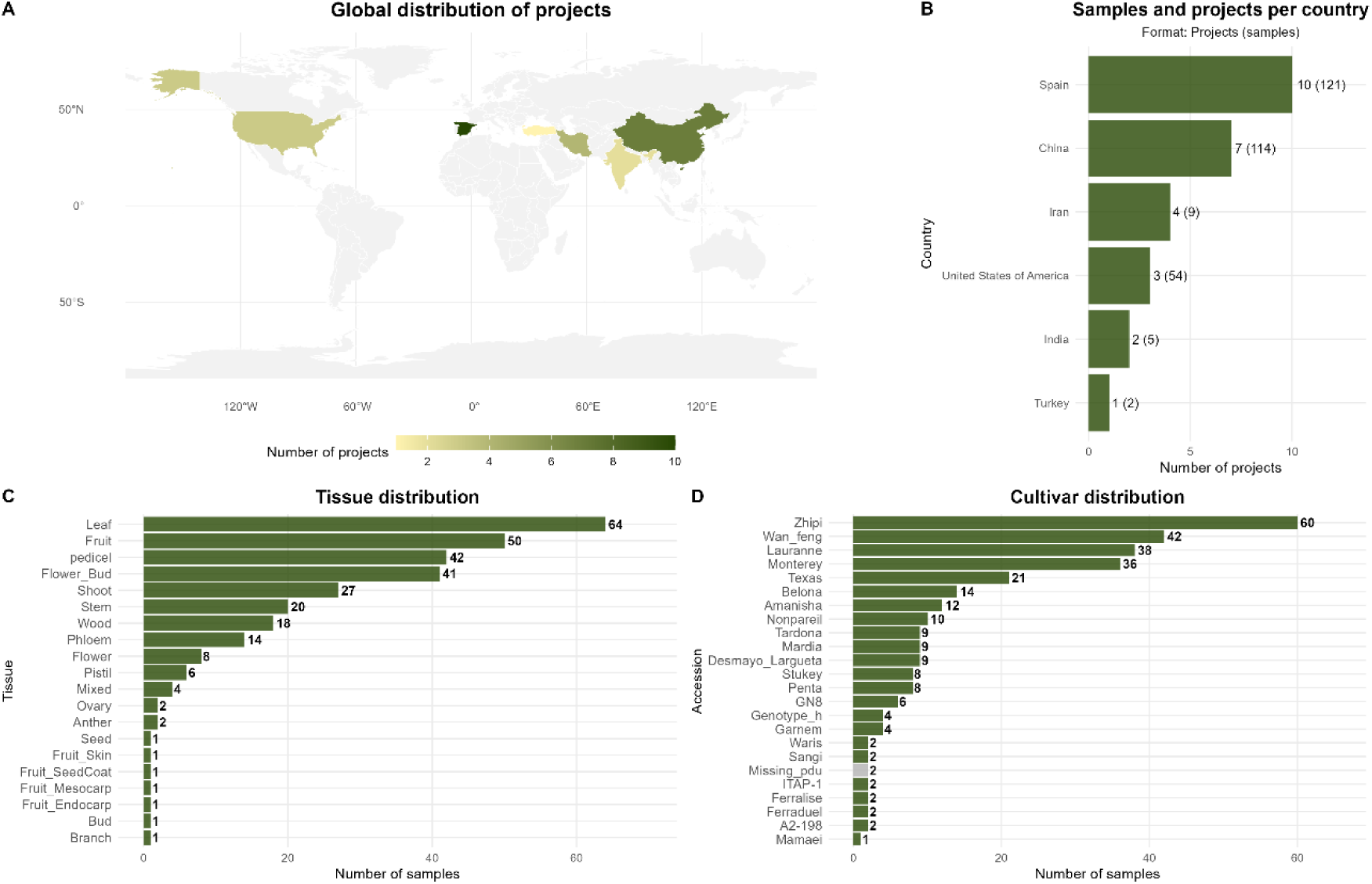
Geographic and biological characterization of 305 RNA-Seq samples from Prunus dulcis available in NCBI SRA. (A) Global distribution of projects. (B) Projects by country and number of samples. (C-D) Sample distribution per tissue and cultivar.

### Expression atlas dataset composition

Following metadata curation, exclusion of low-quality samples (low coverage, low insert size, and pseudoalignment rates below 80%; Supplementary Table S1), and unification of technical replicates (see Materials and Methods), 156 high-quality paired-end samples were retained. Notably, 71% of the projects included a single tissue type, whereas 29% included multiple tissue types. For atlas construction, only untreated or control samples representing baseline expression states were selected. This yielded a final reference dataset of 142 samples, covering 12 distinct tissues from 17 independent projects, for downstream analyses.

The atlas dataset showed a similar tissue distribution, with flower buds (22.5%), pedicel (21.1%), and fruit and leaf (15.5% each) as the most represented tissues. The atlas encompasses 15 cultivars, with ‘Monterey’ (24.6%) and ‘Zhi pi’ (21.1%) being the most represented. The number of samples per project ranged from 1 to 36 (median = 3), reflecting varying study scales and experimental designs.

### Global distribution of samples and tissues across expression datasets

To explore sample-level variability, we visualized the global distribution of samples based on the genes with the highest expression variability using a dimensionality reduction plot with t-SNE (see Materials and Methods) applied to gene-level estimated counts. Samples clustered primarily by tissue type, and tissues belonging to different projects, such as flowers and flower buds, grouped together (Fig. 3). However, a secondary structure by project was also detected, and tissue and project were strongly confounded, as most projects contributed a single tissue. Phloem samples (n = 4) showed higher dispersion than other tissues from the same project, and three leaf samples showed a slight deviation from their group. Quality control metrics for these samples were within the normal range observed across the dataset. Therefore, these deviations may reflect differences between projects or developmental stages.

**Figure 3.**
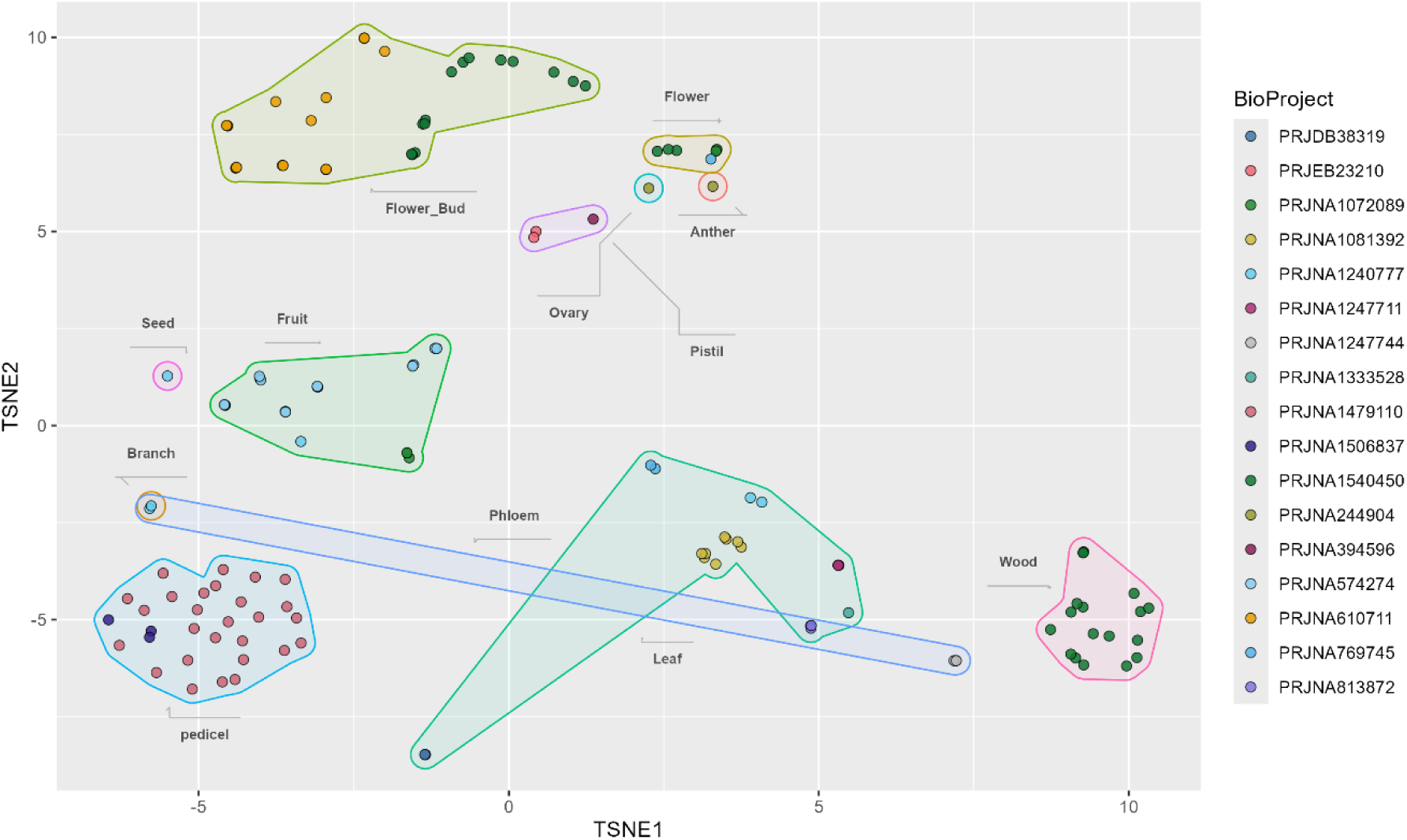
t-SNE visualization showing the global distribution of samples based on gene expression profiles of the 5,000 highly variable genes (HVGs), using a perplexity value of 30.

### Gene composition

The *Prunus dulcis ‘*Texas’ v2.0 genome contains 25,225 annotated genes across the eight chromosomes, of which 18,293 genes (72.52% of the total) showed detectable expression in at least one of the tissues included in this expression atlas. Of these expressed genes, 2,612 (14.28%) were expressed in all tissues. The number of expressed genes (Table 1) varied across tissues, ranging from 8,568 genes in anthers to 14,154 genes in flower buds, with no significant association between the number of expressed genes and sample size per tissue (Spearman’s ρ = 0.13, p = 0.683).

**Table 1.** Summary of expressed and tissue-specific genes across the two methodologies.

| Tissue | Expressed Genes | Expressed % | Bgee-specific Genes | Bgee % | TPM-specific Genes | TPM % | Szymkiewicz-Simpson Coefficient | No. Samples |
|---|---|---|---|---|---|---|---|---|
| **Anther** | 8,568 | 33.97 | 263 | 3.07 | 1,110 | 12.96 | 0.98 | 1 |
| **Branch** | 13,994 | 55.48 | 18 | 0.13 | 687 | 4.91 | 1.00 | 1 |
| **Flower** | 13,539 | 53.67 | 22 | 0.16 | 344 | 2.54 | 0.91 | 7 |
| **Flower bud** | 14,154 | 56.11 | 45 | 0.32 | 280 | 1.98 | 0.69 | 32 |
| **Fruit** | 11,414 | 45.25 | 6 | 0.05 | 82 | 0.72 | 0.50 | 22 |
| **Leaf** | 11,986 | 47.52 | 24 | 0.20 | 520 | 4.34 | 0.83 | 22 |
| **Ovary** | 12,445 | 49.34 | 14 | 0.11 | 273 | 2.19 | 1.00 | 1 |
| **Phloem** | 13,053 | 51.75 | 3 | 0.02 | 170 | 1.30 | 1.00 | 4 |
| **Pistil** | 13,831 | 54.83 | 18 | 0.13 | 188 | 1.36 | 0.72 | 3 |
| **Seed** | 12,540 | 49.71 | 65 | 0.52 | 490 | 3.91 | 0.98 | 1 |
| **Wood** | 11,701 | 46.39 | 13 | 0.11 | 245 | 2.09 | 0.92 | 18 |
| **Pedicel** | 13,354 | 52.94 | 17 | 0.13 | 520 | 3.89 | 1.00 | 30 |

### Tissue-specific genes

The tissue specificity analysis identified 508 tissue-specific genes with the Bgee approach, of which 491 were also detected by the median TPM approach. The median TPM approach additionally identified 4,418 tissue-specific genes not detected by the Bgee approach. To assess the concordance between both methods, we calculated the Szymkiewicz-Simpson coefficient (Fig. 4A). Overlap varied across tissues, with Szymkiewicz-Simpson coefficients ranging from 0.50 to 1.00. The high mean Szymkiewicz-Simpson coefficient of 0.88 indicates that most genes detected by the Bgee approach were also detected by the median TPM approach. Both approaches yielded moderately concordant rankings of tissues by the number of tissue-specific genes detected (Kendall’s τ = 0.55, p = 0.013). We also analyzed the relationship between the number of samples per tissue and the number of tissue-specific genes detected (Fig. 4B). No significant association was observed between sample size and the number of detected tissue-specific genes (TPM Spearman’s ρ = -0.234, p = 0.464; Bgee Spearman’s ρ = -0.18, p = 0.575).

**Figure 4.**
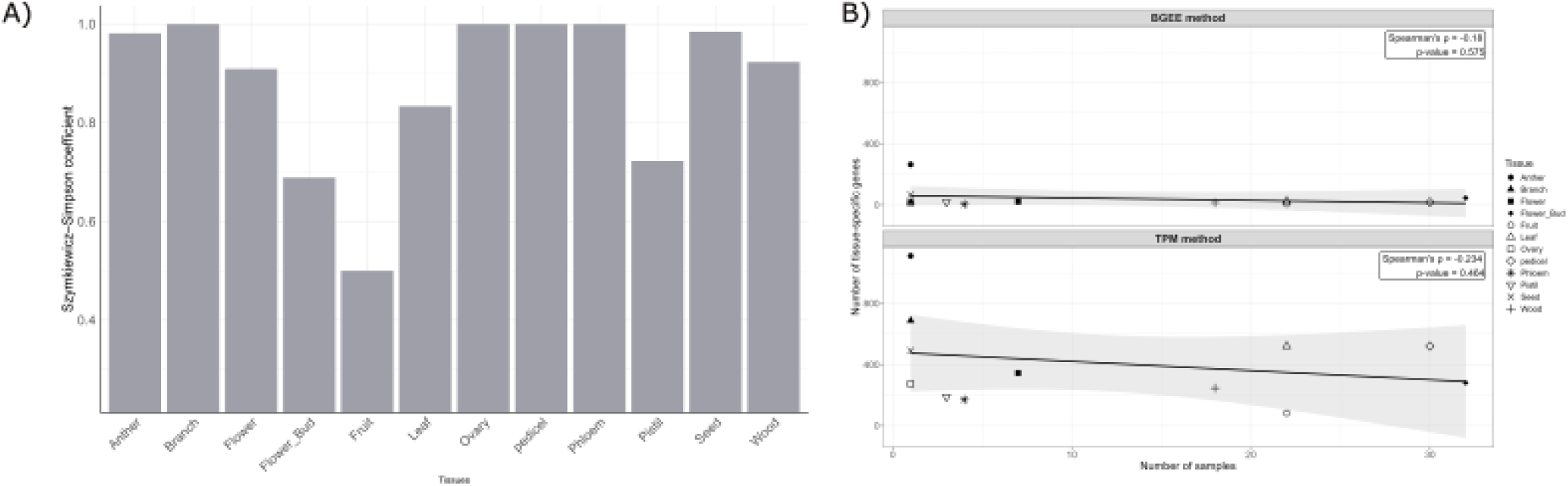
A) Similarity analysis between approaches using the Szymkiewicz-Simpson coefficient. B) Relationship between the number of samples and the number of tissue-specific genes detected.

The distribution of tissue-specific genes was uneven across tissues (Table 1). Anther was consistently the tissue with the highest proportion of tissue-specific genes under both approaches, despite having the lowest total number of expressed genes, suggesting a highly specialized transcriptional program. To further characterize these tissue-specific genes, we performed a functional enrichment analysis. The analysis was carried out primarily using *Arabidopsis thaliana* orthologs (Supplementary Table S4), taking advantage of the more comprehensive functional annotation of this model species. Since a moderate proportion of tissue-specific genes lacked an *Arabidopsis* ortholog (4.4-20.6% depending on the tissue; Table 2), we additionally performed enrichment directly in *Prunus dulcis* using its own functional annotation (Supplementary Table S5). The *Arabidopsis*-based analysis detected significant enrichment in more tissues and generally yielded more enriched GO terms, and both approaches were moderately to highly concordant in the main biological processes, depending on the tissue (mean semantic similarity of 0.60; Table 2). The terms exclusively recovered by STRING corresponded mostly to broad regulatory categories or to more general terms for processes already captured at higher resolution by the *Arabidopsis* enrichment. In a few cases, the *P. dulcis* enrichment additionally recovered specific processes not detected in *Arabidopsis* (e.g., Ethylene-activated signaling pathway in fruits), which were incorporated to complement the characterization.

**Table 2.** Comparison of functional enrichment results obtained using *Arabidopsis* thaliana orthologs and Prunus dulcis annotations for tissue-specific genes detected with median TPM. For each tissue, the table shows the total number of tissue-specific genes, the number and percentage lacking an *Arabidopsis* ortholog, the number of significantly enriched GO Biological Process terms identified using *A. thaliana* orthologs and *P. dulcis* annotations, and the semantic similarity (mgo_Sim) between both enrichment approaches.

| Tissue | Total specific | Genes without *Arabidopsis* ortholog (n) | Genes without *Arabidopsis* ortholog (%) | GO terms *Arabidopsis* | GO terms *P. dulcis* | mgo_Sim |
|---|---|---|---|---|---|---|
| Wood | 245 | 27 | 11.02 | 0 | 10 | - |
| Seed | 490 | 51 | 10.41 | 5 | 2 | 0.50 |
| Branch | 687 | 62 | 9.02 | 61 | 30 | 0.75 |
| Phloem | 170 | 35 | 20.59 | 28 | 3 | 0.44 |
| Pistil | 188 | 34 | 18.09 | 9 | 1 | 1.00 |
| Flower | 344 | 40 | 11.63 | 53 | 4 | 0.40 |
| Fruit | 82 | 9 | 10.98 | 6 | 2 | 0.17 |
| Flower bud | 280 | 30 | 10.71 | 68 | 38 | 0.72 |
| Leaf | 520 | 23 | 4.42 | 86 | 42 | 0.84 |
| Anther | 1110 | 190 | 17.12 | 39 | 16 | 0.61 |
| Ovary | 273 | 42 | 15.38 | 15 | 0 | - |
| Pedicel | 520 | 91 | 17.50 | 20 | 0 | - |

To interpret the functional landscape across different tissues, we constructed a tissue-specific GO semantic network with the top 15 significant GO terms obtained from the enrichment analysis (Supplementary Table S4). This approach effectively reduces semantic redundancy by grouping semantically related terms.

The resulting network (Fig. 5) revealed GO term modules associated with distinct biological processes across tissues. In leaf tissue, the terms were predominantly associated with photosynthesis (GO:0015979) and related processes, including photosynthetic electron transport chain (GO:0009767) and terms related to tetrapyrrole metabolic process (GO:0033013). In branch, clusters were related to cell wall organization (GO:0071555), phenylpropanoid metabolism (GO:0009698), and plant-type secondary cell wall biogenesis (GO:0009834), consistent with structural and lignification functions.

**Figure 5.**
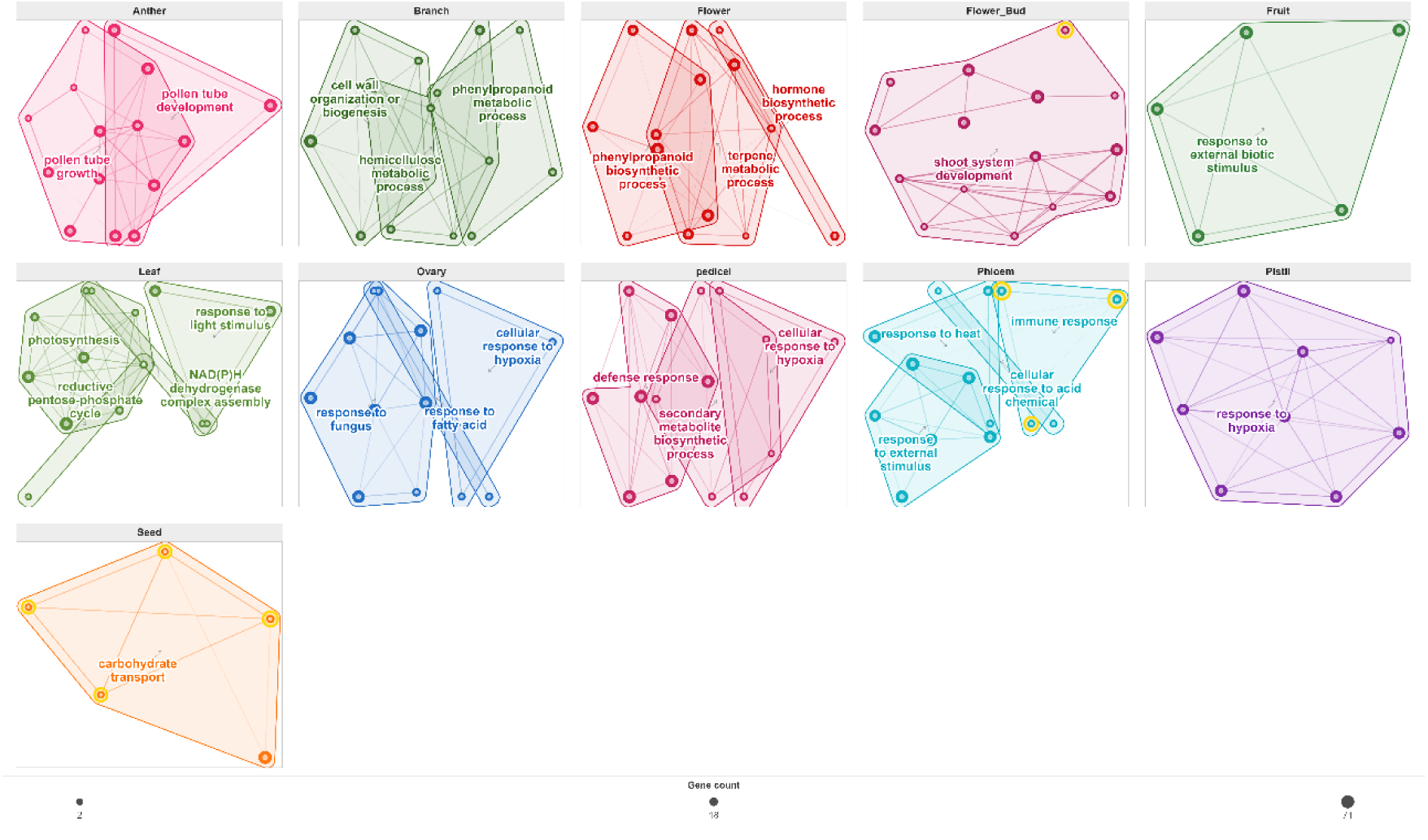
GO semantic similarity networks of the top 15 significant GO terms resulting from the functional enrichment analysis of tissue-specific genes for each tissue. The yellow halo highlights GO terms identified by both the Bgee and the median TPM approaches.

Among reproductive tissues, flower bud GO terms clustered around shoot system development (GO:0048367), including flower development (GO:0009908), and phyllome development (GO:0048827). In anthers, clusters were mainly related to pollen tube growth (GO:0009860), pollen tube development (GO:0048868), and various forms of cell growth (GO:0048588, GO:0009826).

In flowers, the dominant clusters corresponded to secondary metabolism, including phenylpropanoid biosynthetic process (GO:0009699), terpene and terpenoid metabolism (GO:0042214, GO:0006721), and pigment metabolic process (GO:0042440), which are associated with the production of color and scent. Seeds were characterized by clusters involved in carbohydrate transport (GO:0008643), especially sucrose transport (GO:0015770) and transmembrane transport (GO:0055085). Finally, in fruits, the main cluster was associated with response to external biotic stimulus (GO:0043207) and fatty acid beta-oxidation using acyl-CoA oxidase (GO:0033540). The complementary *P. dulcis* functional analysis additionally highlighted the ethylene-activated signaling pathway (GO:0009873) in fruit (Supplementar Table S5). Overall, semantic clustering revealed biologically coherent functional profiles that differed among tissues.

Since several tissues were sampled across multiple developmental stages, tissue-level profiles may dilute stage-specific biological signals and reduce the detection of tissue-specific genes. For this reason, we also evaluated specificity across developmental stages within each tissue, which revealed higher agreement between methods (Kendall’s τ = 0.60, p = 3.387e-6), suggesting that developmental stage is a source of transcriptional variation within tissues in our dataset. Functional enrichment of developmental-stage-specific genes (Supplementary Table S6) revealed significantly enriched biological processes in developmental stages that showed no significant enrichment when analyzed at the tissue level. For example, fruit samples displayed enrichment for cell division-related processes, such as sister chromatid segregation (GO:0000819), and for defense-related processes (GO:0006952) at 12 DAF, whereas later stages were characterized by ethylene signaling (GO:0009873) and cellular response to hypoxia (GO:0071456). Similarly, flower buds transitioned from cytokinin catabolism and lipid/secondary metabolism (GO:0009823, GO:0016114) in early stages to floral organ development and pollen wall assembly (GO:0048437, GO:0010208) at intermediate stages, followed by fatty acid biosynthesis and phenylpropanoid metabolism (GO:0006636, GO:0009699) in the final stages, thus providing substantially higher functional resolution.

### Expression Atlas Tools

FruitDBase tools can be accessed from both the toolbar and the homepage. The homepage includes key performance indicators that summarize the data available for each species (Fig. 6A), as well as links to relevant resources for each species (Fig. 6B). The tools included in the first version of FruitDBase are as follows:

I. **Search Engine**: Users can search for *Prunus* samples and projects available in the database. Filters allow users to identify specific datasets and explore their metadata, with direct links to the original sources to ensure traceability (Fig. 6C-D).
II. **Expression Explorer**: Users can explore gene expression in two ways: (i) searching for one or multiple genes and (ii) filtering for tissue-specific or developmental-stage-specific genes. In the first mode, users can search for one or multiple genes of interest by name, whereas in the second mode, they can filter genes according to tissue or developmental stage and specify the number of genes to display. In both modes, expression data can be explored with either of two complementary approaches: the Bgee approach or the median TPM approach.

**Figure 6.**
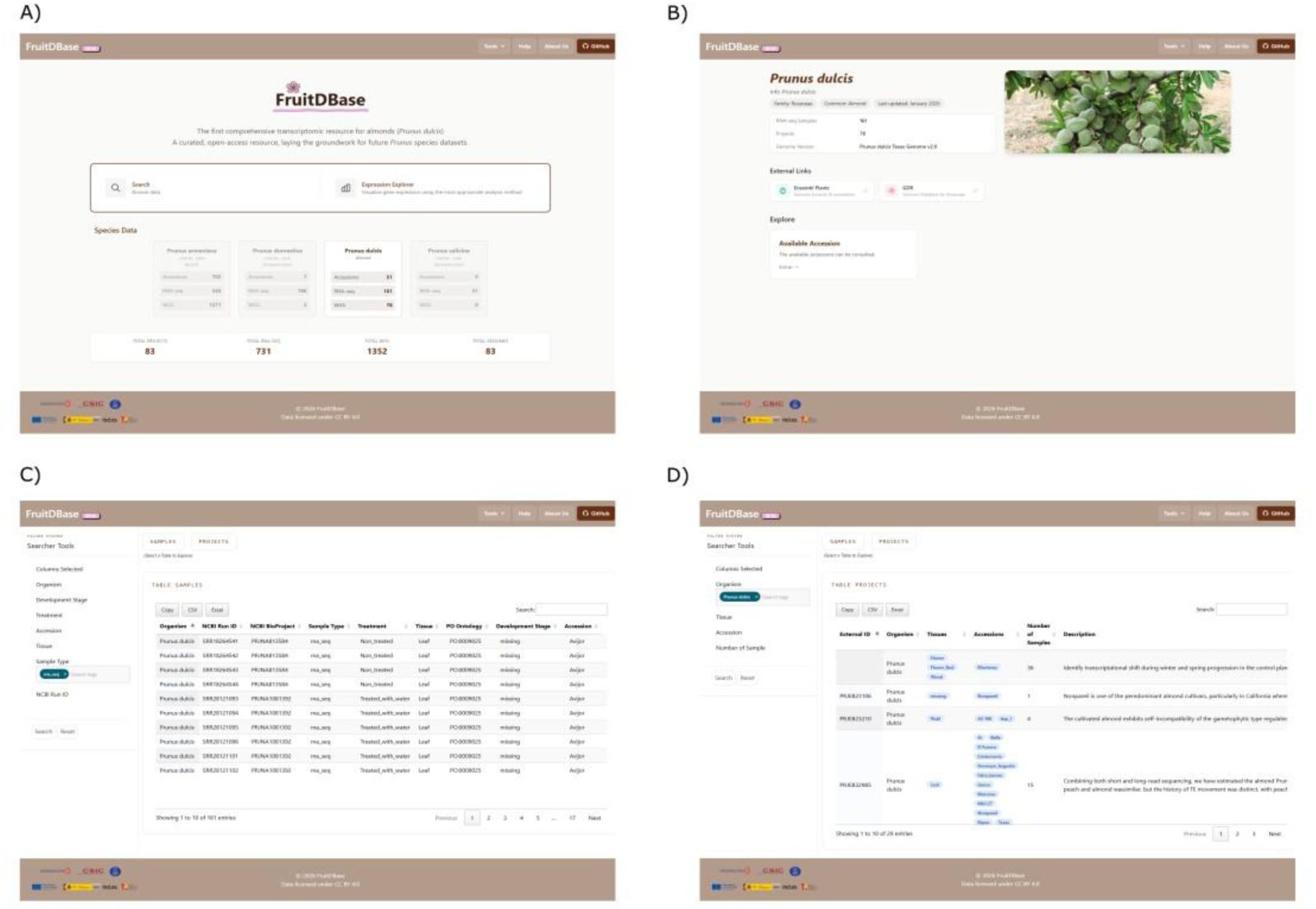
FruitDBase homepage and Search Engine tool. A) Homepage with species-level key performance indicators. B) Species information page. C) Sample search table. D) Project search table.

For the Bgee approach (Fig. 7A), the interface includes five visualization components: (1) a heatmap, which allows exploration and comparison of gene expression scores by tissue or developmental stage. When a gene is selected, additional visualizations become available: (2) a gene information card, containing details such as genomic location and associated GO terms; (3) a bar plot, providing an easy way to compare tissue expression; (4) an expression data table, presenting the same data as the bar plot in tabular format; and (5) a detailed sample-level view: when a tissue is selected in the expression data table, an additional table displays the metadata of the samples used to compute the expression score in that tissue (e.g., developmental stage, external sample ID, TPM value, and presence/absence p-value). In the median TPM approach (Fig. 7B), three visualizations are available: (1) a table with gene information, containing the same details as the gene information card described above; (2) a t-SNE plot to explore expression in context, for example, whether it follows a pattern by tissue or by another grouping variable; and (3) bar plots for comparing expression by tissue or developmental stage.

**Figure 7.**
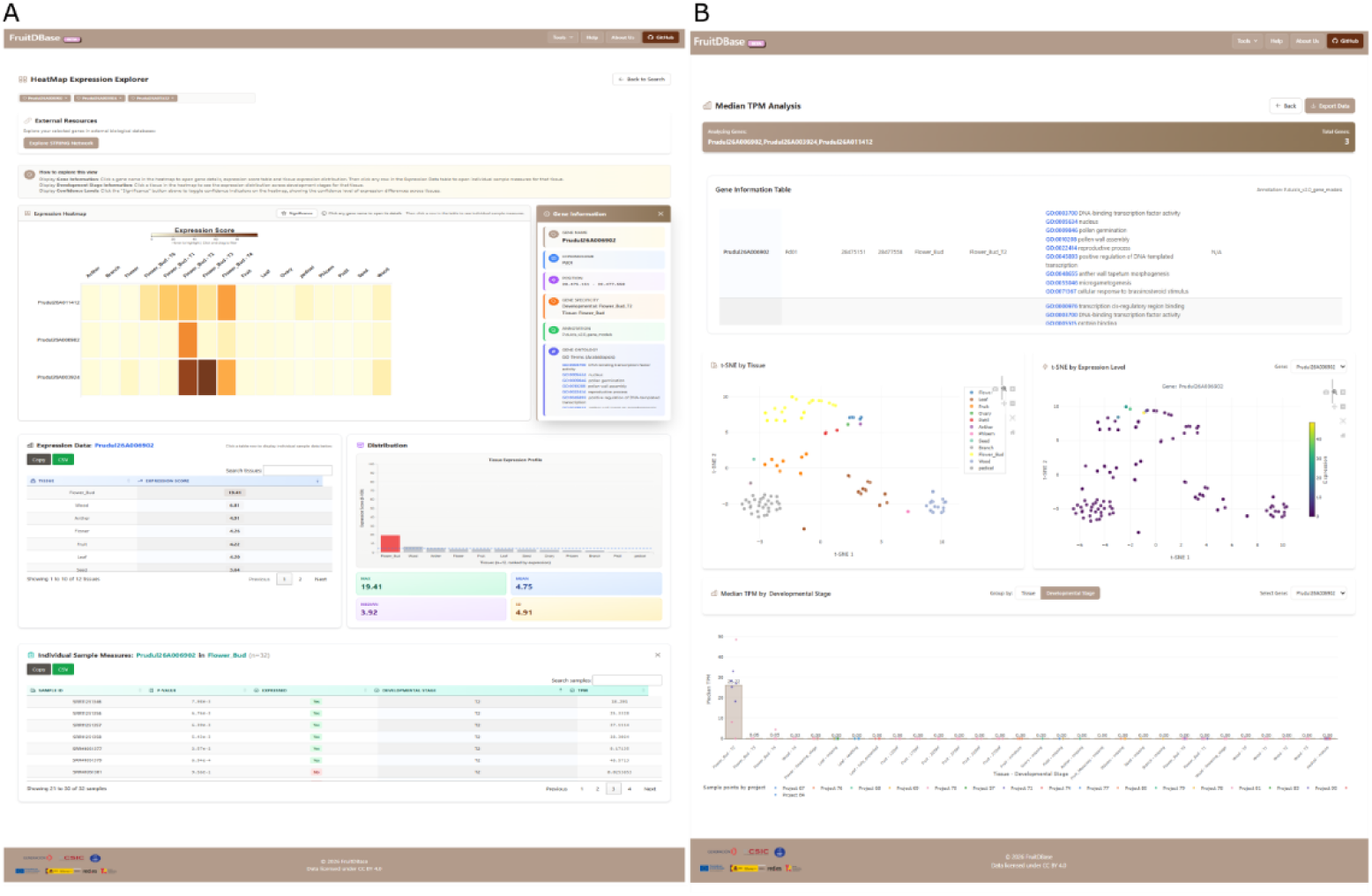
Expression Explorer Tool. A) Bgee approach interface. B) Median TPM approach interface.

### Use Cases

To demonstrate the utility and reliability of our tool, we present two use cases. The first illustrates the use of the web interface for directly exploring and retrieving biological information, whereas the second demonstrates how the generated datasets can be downloaded from the web and further analyzed locally.

#### Use Case I - Using the web interface: Tissue-specific genes involved in endodormancy to ecodormancy release

Bud dormancy in *Prunus* species comprises three physiological stages: paradormancy, endodormancy, and ecodormancy, each associated with distinct transcriptional programs. The transition from endodormancy (stage T1) to ecodormancy (stage T2) represents a critical window, as endodormancy release largely determines flowering time. Therefore, identifying molecular markers of this transition has direct implications for almond breeding programs in the context of climate adaptation [43,45].

As a first case study, we asked whether the expression atlas could identify genes that were highly specific to the ecodormancy stage. Using the search-by-filter option of the Expression Explorer tool available in FruitDBase, we retrieved genes highly specific to flower buds at stage T2. We then explored their expression patterns using the Bgee approach and analyzed their Gene Ontology (GO) enrichment with STRING, accessed via the external resource button. GO enrichment analysis revealed that most of the significantly enriched terms were directly or indirectly associated with pollen development, such as pollen wall assembly (GO:0010208), pollen exine formation (GO:0010584), or anther development (GO:0048653). This result is consistent with previous studies: microsporogenesis has been associated with the ecodormancy phase in almond [46], and pollen development genes have been proposed as transcriptional markers of the endodormancy to ecodormancy transition [43]. To further assess the consistency of our results with previous findings, we examined the expression patterns of the candidate EcoGS genes proposed by Prudencio et al. [43]. To this end, we searched for each gene by name and, using the Median TPM visualization, inspected the tissue- and developmental stage-specificity fields in the gene information panel. Of the seven EcoGS candidate genes, four were classified as highly specific to flower bud T2 samples (Table 3).

**Table 3.** Validation of candidate EcoGS genes from Prudencio et al. [43] using tissue-specific gene analysis.

| **Eco Cluster Genes Candidates (Prudencio)** | **Gene ID (Texas Genome Reference)** | **Gene annotation** | **Match** |
|---|---|---|---|
| Prudu.01G238900 | Prudul26A017732 | GLUCAN ENDO-β-1,3-GLUCOSIDASE | No |
| Prudu.05G071900 | Prudul26A025760 | ABI3 INTERACTING PROTEIN 2, E3 LIGASE | Yes |
| Prudu.05G129900 | Prudul26A007897 | ENDO-β-1,4-GLUCANASE | No |
| Prudu.05G142100 | Prudul26A004276 | BIDIRECTIONAL SUGAR TRANSPORTER SWEET10 | No |
| Prudu.08G187400 | Prudul26A015775 | XYLOGLUCAN ENDOTRANSGLUCOSYLASE 2 | Yes |
| Prudu.08G242500 | Prudul26A021485 | 4-COUMARATE:CoA LIGASE-LIKE 1 | Yes |
| Prudu.01G134200 | Prudul26A010948 | AQUAPORIN, NODULIN-LIKE INTRINSIC PROTEIN 7 | Yes |

#### Use Case II - Repository data download and analysis: Selection of stably expressed reference genes in almond

In gene expression analyses such as RT-qPCR, normalisation with reference genes whose expression remains stable across samples is essential [47]. Czechowski et al. [44] used a large collection of *Arabidopsis* transcriptomic data to identify novel reference genes that were more stable than traditional ones and suggested that their orthologs could serve the same purpose in other species. Following this approach, we used the atlas to evaluate these candidate genes in almond.

Of the 17 novel reference genes described by Czechowski et al. [44], 13 had an expressed putative ortholog in the atlas (UBC9 through its paralog UBC10), one had a putative ortholog that was not expressed in any tissue (AT4G38070), and three (AT1G13320, AT1G58050 and AT1G62930) were not the best hit of any almond gene and could not be assessed. Of the five traditional reference genes, four could be evaluated (UBQ10, EF-1α, UBC21 and GAPDH), EF-1α through paralogs detected by the same microarray probe set [44], whereas ACT2 had no putative ortholog (Supplementary Table S7). Nine of the 13 novel genes and two of the four traditional genes (UBQ10 and EF-1α) were supported in almond, ranking within the 25% most stable genes of the atlas (Fig. 8). SAND (AT2G28390) and TIP41-like (AT4G34270) were the most stable candidates, ranking within the 3.5% and 6.5% most stable genes, respectively. Stability often differed between orthologs of the same Arabidopsis gene, as observed for the copies of UBQ10, EF-1α and AT4G33380.

**Figure 8.**
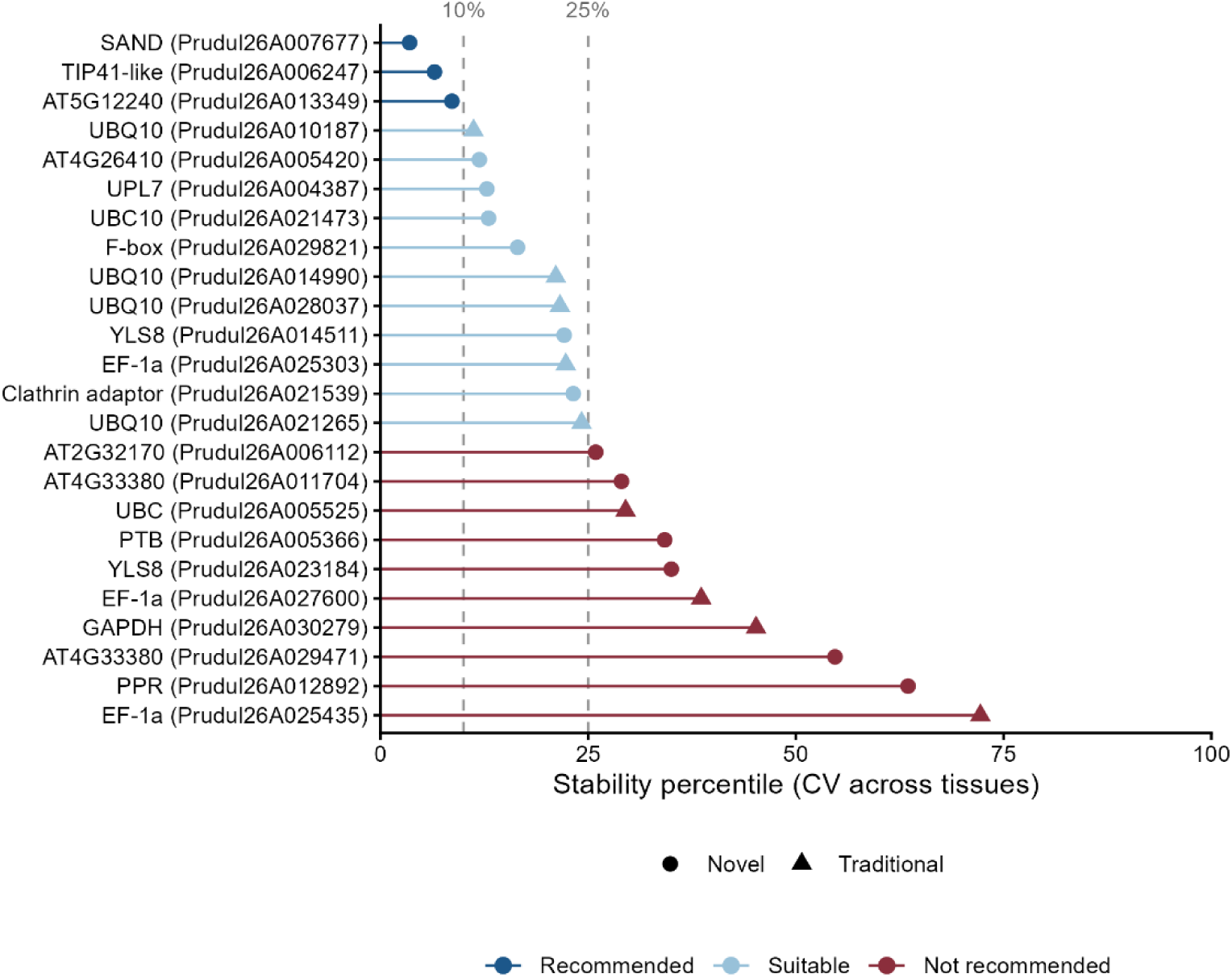
Expression stability in almond of putative orthologs of Arabidopsis reference genes. Stability percentile of the CV across tissues for each almond gene. Dashed lines mark the 10th and 25th percentiles. Shapes distinguish novel (circles) and traditional (triangles) reference genes described by Czechowski et al. [44]

## Discussion

FruitDBase provides a common platform to explore publicly available transcriptomic data for *Prunus dulcis* in the NCBI SRA, unifying previously scattered datasets in a single place and providing access to standardized metadata. The database is updated periodically to maintain and expand its data content. In addition, through an accessible and easy-to-use interface, expression can be explored using two approaches, allowing users to choose the methodology best suited to their specific needs.

Our results highlight differences in tissue specificity detection between the two approaches in heterogeneous datasets. The median TPM approach detects a substantially higher number of tissue-specific genes, which may reflect greater sensitivity. However, due to the absence of cross-study normalization, this broader detection is also more susceptible to technical variability between projects.

In contrast, the Bgee approach provides a more conservative set of tissue-specific genes. Being rank-based, it is less sensitive to differences in scale between studies [31], making it better suited for comparing samples from different projects, at the cost of detecting fewer tissue-specific genes.

The comparison between the two approaches further supports this interpretation. Despite the substantial differences in the total number of detected genes, the high Szymkiewicz-Simpson coefficient indicates that most genes identified by the Bgee approach were also recovered by the median TPM approach. This indicates that the Bgee approach captures a core set of highly specific genes, whereas the TPM approach extends this set by including genes with moderate or variable expression patterns. Thus, the choice of approach should depend on the specific research objectives. The median TPM approach may be more suitable for exploratory analyses and candidate gene discovery, whereas the Bgee approach, designed to improve comparability across studies, is better suited for cross-study comparisons and for analyses that prioritize a more conservative set of candidates.

The two use cases presented in this study support both the biological coherence and the practical utility of the atlas. In the dormancy use case, the recovery of pollen development genes as markers of the endodormancy to ecodormancy transition was consistent with previously published findings [43]. In addition, in the reference gene use case, the atlas showed its utility for prioritising candidate genes, by selecting the most suitable reference genes from a list of candidates previously described in *Arabidopsis*. SAND and TIP41-like, two of the most stable novel reference genes in *Arabidopsis* [44], were also among the most stable genes across almond tissues. This may facilitate gene expression studies in almond, where reference genes are commonly adopted from peach or other species and validated in each study [48,49], since candidate genes can first be evaluated in the atlas before being tested experimentally, thereby reducing the number of genes that require RT-qPCR validation.

The present version of the atlas has some limitations. The limited sample size available for certain tissues, such as anther and ovary, may affect the representativeness of their global expression profiles. Furthermore, for tissue-level expression, heterogeneous samples across developmental stages were integrated. This approach may result in more diffuse global expression signals in cases where transcriptional profiles differ substantially between developmental stages. We addressed this by providing data at both the tissue and developmental-stage levels, although sample sizes are smaller for the latter. In any case, the repository is designed to integrate new samples, both for tissues already represented in the atlas and for new tissues, as soon as they become available.

The influence of cultivar-specific differences on gene expression was not evaluated in this study. Moreover, cultivar representation was not homogeneous across tissues: while tissues such as leaf and flower included balanced contributions from multiple cultivars, others were represented by a single cultivar, such as fruit with ‘Zhi pi’, or were predominantly represented by one cultivar, such as flower buds with ‘Monterey’. Consequently, tissue-specific expression patterns identified for these particular tissues should be interpreted with additional caution, as cultivar-specific effects cannot be fully disentangled from tissue-specific ones. Accordingly, users should interpret expression profiles as reflecting a general *Prunus dulcis* transcriptome rather than cultivar-specific patterns, bearing in mind that this generalization is more robust for tissues with balanced cultivar representation. Ideally, expression profiles would be resolved by cultivar, developmental stage, and tissue type, but this was not feasible given the current availability of data.

In addition, the confounding between tissue and project highlights the need for new experiments in which the same tissues are sampled by multiple independent projects, as well as for projects covering several tissues, which would allow differences in expression between tissues to be better resolved. Until then, tissue-level expression patterns should be interpreted with the possible influence of technical effects in mind.

Because FruitDBase depends on the availability and quality of public datasets, certain tissues or developmental stages remain underrepresented. This underrepresentation may be partly attributable to the technical challenges associated with sampling and RNA extraction from some specialized tissues, as well as to differences in research priorities. By contrast, tissues such as flowers or fruits are more frequently studied because of their direct relevance to fruit production, development and quality. Nevertheless, the current atlas provides a comprehensive representation of gene expression based on the transcriptomic data publicly available to date. As new transcriptomic data become available, the coverage and resolution of this database are expected to improve.

In summary, the expression atlas hosted in FruitDBase represents a practical and accessible resource that supports both exploratory and hypothesis-driven research, contributing to functional genomics studies and the identification of candidate genes in *Prunus dulcis*.

## Conclusion

The Almond Expression Atlas v1.0 constitutes the first comprehensive transcriptomic resource for *Prunus dulcis*, integrating publicly available RNA-Seq data into a unified and standardized platform. By combining two complementary expression approaches, the atlas allows researchers to explore gene expression patterns across tissues and developmental stages using the approach best suited to their needs, from broad exploratory searches to more conservative cross-study comparisons. This resource represents the first tool implemented within FruitDBase, laying the foundation for future transcriptomic and genomic tools to be developed for *Prunus* species. As new datasets become available, FruitDBase is expected to keep expanding, consolidating its role as a centralized, reusable repository for the *Prunus* research community.

## Data availability

The raw RNA-Seq data analyzed in this study are publicly available in the Sequence Read Archive (SRA). The accession numbers for all samples are listed in Supplementary Table S3. Data from the unpublished UC Davis project (PRJNA1540450) are under embargo and will be publicly released upon publication of the associated study.

## Availability of Source Code and Requirements

Project name: FruitDBase

Project home page: https://github.com/evalopezf/FruitDBase

Operating system(s): Platform independent

Programming language: R, Shell, Python (pipeline); PHP, MySQL, JavaScript (web application)

License: GNU GPL v3

RRID: SCR_029077

bio.tools ID: fruitdbase

## List of abbreviations

bp: Base pairs
DAF: Days After Flowering
ENA: European Nucleotide Archive
FAIR: Findable, Accessible, Interoperable and Reusable
FDR: False Discovery Rate
GDR: Genome Database for Rosaceae
GO: Gene Ontology
HVGs: Highly Variable Genes
RNA-Seq: RNA Sequencing
SRA: Sequence Read Archive
t-SNE: t-distributed Stochastic Neighbor Embedding
TPM: Transcripts Per Million

## Additional Files

**Supplementary Material S1.** Detailed explanation of metadata curation.

**Supplementary Figure S1.** Histogram showing the distribution of insert size peak values.

**Supplementary Table S1.** Summary of sequencing quality control and pseudoalignment statistics for all RNA-Seq samples.

**Supplementary Table S2.** Global almond production by country according to FAOSTAT, 2023.

**Supplementary Table S3.** Curated metadata of all publicly available Prunus dulcis RNA-Seq samples retrieved from the Sequence Read Archive (SRA) and used for atlas construction.

**Supplementary Table S4.** Functional enrichment analysis of tissue-specific genes using *Arabidopsis thaliana* orthologs.

**Supplementary Table S5.** Functional enrichment analysis of tissue-specific genes using *Prunus dulcis* annotations.

**Supplementary Table S6.** Functional enrichment analysis of developmental stage-specific genes.

**Supplementary Table S7.** Almond putative orthologs of the *Arabidopsis* reference genes validated by qRT-PCR in Czechowski et al. [44] and their expression stability in the atlas.

## Authors’ contributions

E.M.L-F. curated the data, performed the analyses, and wrote the original draft of the manuscript. M.A.Z. and A.T. provided data and reviewed and edited the manuscript. J.A.B. conceived the study, contributed to the methodology, supervised the work, and edited the manuscript. P.J.M-G. conceived and supervised the study, acquired funding, and wrote and edited the manuscript. All authors read and approved the final manuscript.

## Acknowledgments

P.J.M-G. gratefully acknowledges the support of grant 22838/FLB/24, provided by Fulbright– Fundación Séneca, Región de Murcia (Spain).

E.M.L-F. acknowledges the support of the Momentum CSIC Programme: Develop Your Digital Talent.

## Supplementary Material

### Supplementary Material 1. Detailed explanation of metadata curation

The first step was the creation of a developmental stage column, since the tissue column contained information about the sample’s developmental stage (e.g., Flower_pink_stage). The second step involved unifying and standardizing names in the tissue, treatment, and cultivar columns (e.g., “fruti” and “fruitlets” were changed to “Fruit”). Then, a new nomenclature “Mainpart_Secondpart” was applied to the tissue column whenever the subpart was mentioned in the sample name (e.g. “Fruit_Mesocarp”, “Fruit_SeedCoat”). Finally, missing values were manually completed by searching original sources and publications associated with the corresponding projects, for the treatment, tissue, developmental stage, and cultivar columns.

After manual curation was completed, rows with missing data in essential columns (tissue and treatment) were deleted. Location metadata were also supplemented to characterize the geographical distribution of the datasets. In addition, some projects were excluded from the atlas based on their experimental design, potential redundancy, sequencing strategy, or biological context. The excluded projects were:

- *PRJNA854307:* Whole-genome sequencing rather than RNA-Seq
- *PRJNA813584:* Duplicated project of PRJNA1081392.
- *PRJNA1253203:* Virome analysis in symptomatic almond plants; excluded because the samples represent a disease-associated condition and were not suitable for characterizing baseline gene expression.
- *PRJEB47633:* Grafted plants; excluded because the study investigates the effect of rootstock genotype on scion gene expression.
- *PRJEB50411:* Grafted plants; excluded because the study focuses on scion-rootstock interaction and its effect on gene expression.

**Supplementary Figure 1.**
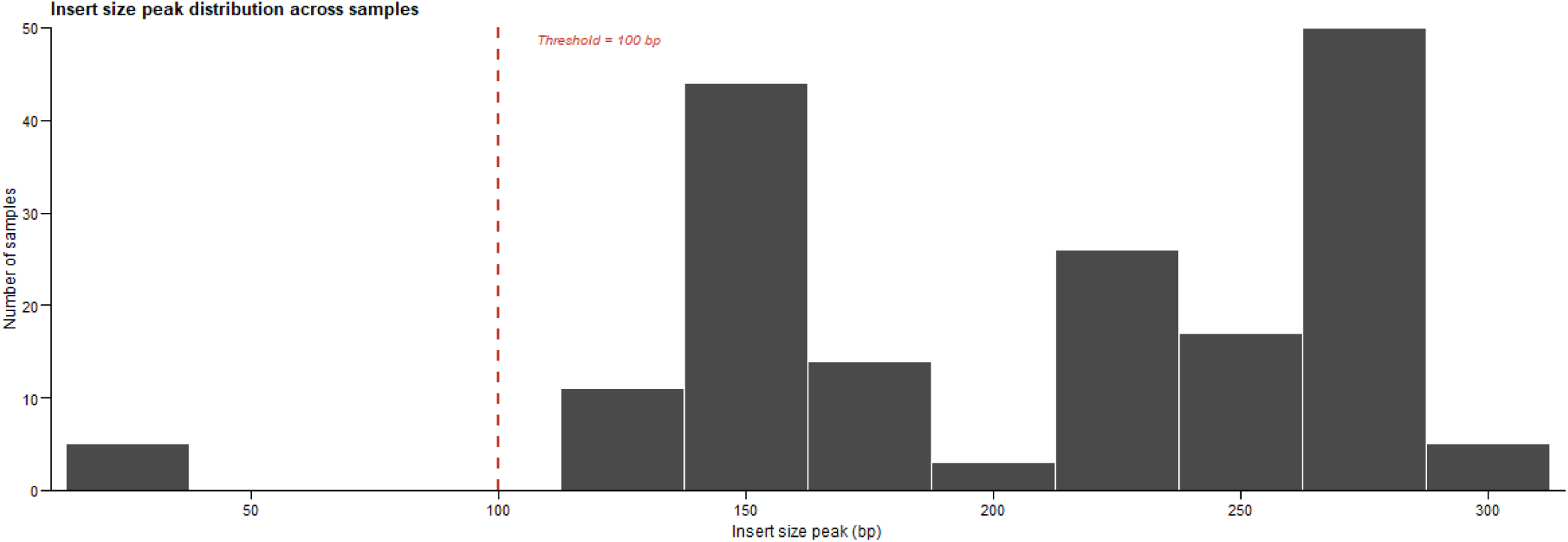
Histogram showing the distribution of insert size peak values (bp) estimated by FastP across all samples prior to filtering. The dashed red line indicates the 100 bp thresholds used to exclude low-quality samples.

